# Functional Validation of Known *BEST1* Variants Using a Robust Halide-Sensitive YFP Quenching Assay

**DOI:** 10.64898/2026.10.19.760219

**Authors:** Sachini U. Siriwardena, Mayur Choudhary, Jason Lee, Benjamin R. Yerxa, Sally Tucker, George Magrath, Devin S. McDougald, Christopher H. Tipper, Ashwath Jayagopal

## Abstract

Bestrophinopathies comprise a heterogeneous group of inherited retinal degenerative diseases caused by mutations in the Bestrophin-1 (*BEST1*) gene. Although more than 250 pathogenic *BEST1* variants have been reported, only a small subset has undergone functional interrogation. OPGx-BEST1 is an investigational gene therapy designed to restore BEST1 function through augmented wild-type BEST1 expression in retinal pigment epithelium (RPE) cells. Previous studies in animal models and patient-derived cellular systems have demonstrated that loss-of-function (LOF) *BEST1* variants can be complemented with wild-type protein expression and are therefore amenable to gene augmentation strategies. In contrast, known gain-of-function (GOF) mutations appear to retain pathogenic activity despite augmentation with wild-type BEST1. Therefore, given the therapeutic relevance of LOF/GOF disease mechanisms in bestrophinopathies, a comprehensive genotype-phenotype characterization across the BEST1 mutational landscape is critical to inform clinical trial enrollment. Conventional approaches to profile *BEST1* variants have relied upon patient-derived RPE models and electrophysiologic assessment of channel function. While these systems provide valuable translational insight, they lack the scalability, speed, and experimental flexibility needed for a systematic evaluation of many variants for patient selection. Here, we describe a functional screening platform that leverages an engineered halide-sensitive reporter cell line to quantitatively assess the activity of BEST1 mutant proteins.

## Introduction

Bestrophinopathies are a group of inherited retinal degenerations caused by pathogenic variants in the *BEST1* gene, which encodes Bestrophin-1, a calcium-activated anion channel predominantly expressed on the basolateral membrane of retinal pigment epithelium (RPE) cells [1]. BEST1 performs a critical role in maintaining retinal homeostasis through the regulation of ion transport, fluid balance, and photoreceptor-RPE interactions. Mutations in BEST1 give rise to a clinically and genetically heterogeneous spectrum of diseases, including Best vitelliform macular dystrophy (BVMD), autosomal recessive bestrophinopathy (ARB), and adult-onset vitelliform macular dystrophy (AVMD) [1, 2]. To date, more than 250 disease-associated variants have been reported across the *BEST1* coding sequence, highlighting the extensive allelic diversity that characterizes this disease family [3].

Despite the substantial progress in defining the genetic architecture of bestrophinopathies, the functional consequences of most mutations remain poorly understood. Historically, pathogenic variants have been categorized according to their effects on protein expression, trafficking, or chloride conductance with most mutants exhibiting loss-of-function (LOF) behavior [4-6]. These observations provide a strong rationale for gene augmentation therapy aimed at restoring normal BEST1 activity through delivery of a wild-type transgene. Moreover, preclinical studies in animal models and patient-derived in vitro systems demonstrate LOF variants can be effectively complemented by exogenous wild-type BEST1 expression [7-12]. In contrast, certain BEST1 mutations are known to exert gain-of-function (GOF) mechanisms that may be refractory to conventional replacement strategies despite wild-type protein augmentation [13-15].

As new therapeutic approaches emerge, understanding the mechanistic basis of individual *BEST1* variants is becoming increasingly important for patient stratification, clinical trial enrollment, and interpretation of treatment outcomes. Consequently, there is a growing need for systematic approaches capable of distinguishing augmentation-responsive variants from those likely to require an alternative therapeutic modality.

Traditional methods used to characterize BEST1 function utilize electrophysiologic measurements in heterologous expression systems or patient-specific iPSC-derived RPE cells [6, 12, 16]. While these approaches provide valuable mechanistic and translational insights, they are labor-intensive, and difficult to scale across the large landscape of *BEST1* variants. The limited throughput of these platforms represents a significant barrier to comprehensive genotype-phenotype mapping and timely clinical decision-making.

To address this challenge, we developed a functional screening platform that leverages a halide-sensitive reporter cell line that quantitatively assesses BEST1 channel activity. Here, we describe the development, validation, and application of this assay to characterize a subset of BEST1 mutants to support mechanism-based patient selection for an investigational gene replacement therapy.

## Results

### Generation of a stable cell line incorporating a halide-responsive fluorescent sensor

To establish a robust platform for functional assessment of BEST1 channel activity, HEK293 cells were engineered to constitutively express the halide-sensitive fluorescent reporter EYFP-H148Q/I152L. This enhanced yellow fluorescent protein (EYFP) variant contains H148Q and I152L substitutions that increase its sensitivity to quenching by halide ions, particularly iodide, which quenches fluorescence more efficiently than chloride [17]. As a result, intracellular accumulation of iodide induces rapid quenching of EYFP fluorescence, enabling real-time monitoring of BEST1-mediated halide transport.

Stable, polyclonal, integration of the EYFP-H148Q/I152L reporter was achieved to ensure consistent expression across experiments and to minimize variability associated with transient transfection. Following selection and expansion, stable reporter expression in HEK293 cells was confirmed by fluorescence imaging (Supplementary Fig. S1). The resulting cell line provided a robust and reproducible fluorescent signal suitable for YFP quenching assays.

### Development and validation of a halide-sensitive YFP quenching assay using *BEST1* variants with known functional phenotypes

To determine whether the engineered cell line could be used to monitor BEST1 channel function, a halide-sensitive EYFP-H148Q/I152L quenching assay was developed (Fig. 1). Stable HEK293 reporter cells were transiently transfected with wild-type (WT) BEST1, the LOF variant R218H [11, 15], the GOF variant Y236C [14, 15], or a neutral stuffer control. BEST1 channel activity was determined by monitoring the intensity of YFP fluorescence over time following an increase in the extracellular concentration of the halide iodide (Fig. 2A, Supplementary Fig. S2).

**Fig. 1.**
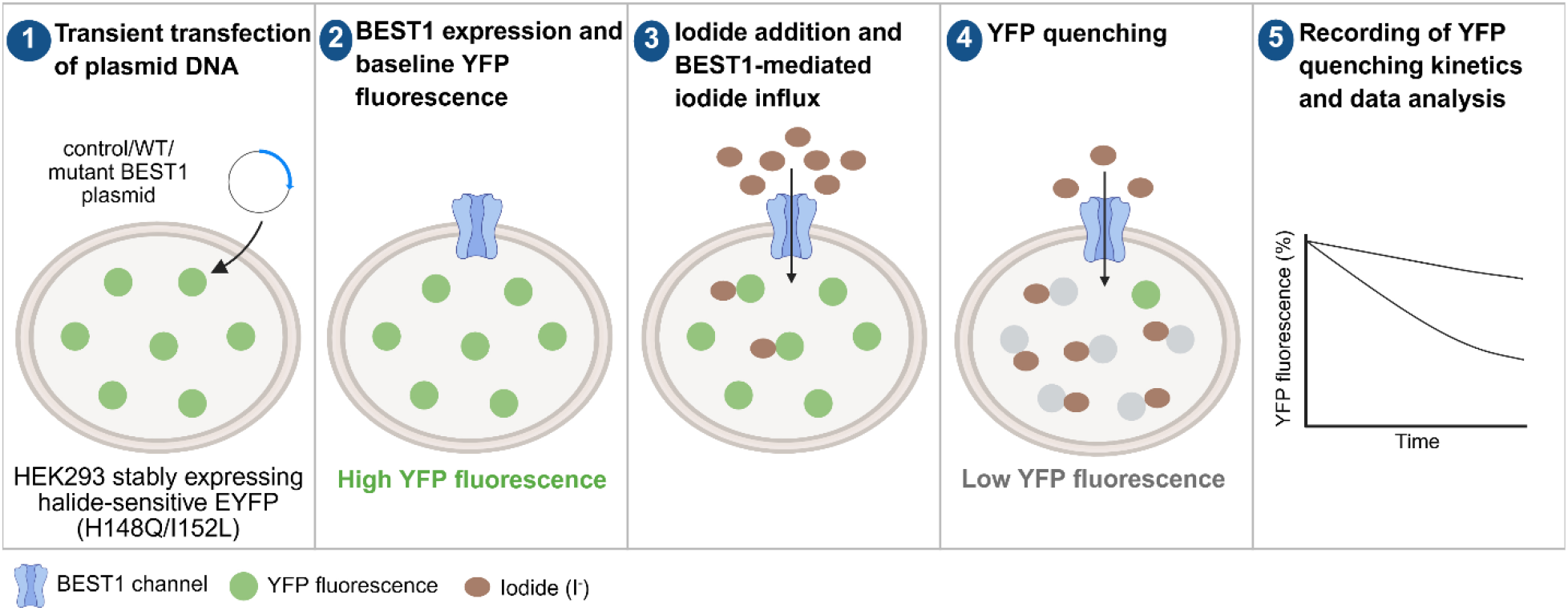
Principle and workflow of the halide-sensitive YFP quenching assay. HEK293 cells stably expressing EYFP-H148Q/I152L were transiently transfected with control, WT or mutant BEST1 plasmids. 48 hours post-transfection, baseline YFP fluorescence was measured. Upon addition of extracellular iodide, BEST1-mediated iodide influx increases intracellular iodide levels, resulting in quenching of the halide-sensitive YFP fluorescence. Fluorescence was recorded over time, and quenching kinetics were analyzed to quantify and compare the functional activity of WT and mutant BEST1 proteins.

**Fig. 2.**
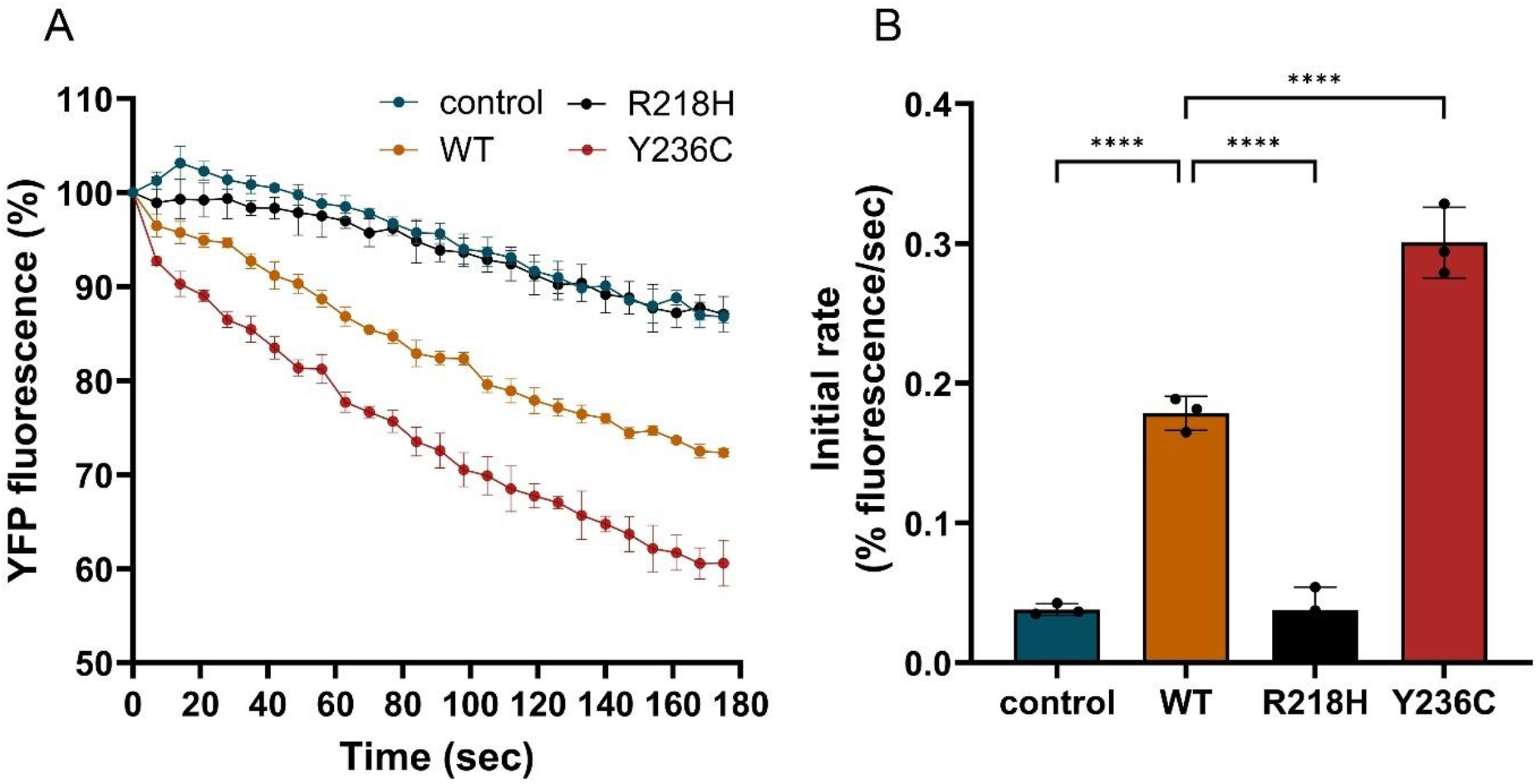
Validation of the halide-sensitive YFP quenching assay using BEST1 variants with known gain- and loss-of-function phenotypes. (A) Representative YFP fluorescence quenching kinetics in HEK293 cells stably expressing halide-sensitive EYFP-H148Q/I152L following transfection with control, WT BEST1, the loss-of-function variant (LOF) R218H, or the gain-of-function variant (GOF) Y236C. Extracellular iodide was added at time 0, and changes in YFP fluorescence were monitored at 7 second intervals for 3 minutes. Fluorescence intensities were normalized to the initial fluorescence signal to calculate % YFP fluorescence. (B) Initial YFP quenching rates calculated from the fluorescence quenching kinetics shown in (A) using the first 60 seconds following iodide addition. Data are presented as mean ± SD (n = 3 technical replicates). Statistical significance was assessed using one-way ANOVA followed by Dunnett’s multiple comparisons test with WT as the reference group. ****P < 0.0001.

The neutral stuffer control revealed a baseline flux of iodide across the plasma membrane as revealed by the slow decay of YFP fluorescence over the assessed time period. This activity was mediated by mechanisms native to HEK293 cells. Ectopic WT BEST1 expression resulted in rapid fluorescence quenching, indicating efficient iodide transport across the plasma membrane. In comparison, mutant BEST1 constructs exhibited distinct quenching profiles, characterized by altered rates of fluorescence decay. The R218H mutant demonstrated a kinetic profile that was largely indistinguishable from the neutral stuffer control, supporting its classification as a LOF variant. In contrast, the Y236C mutant displayed significantly enhanced fluorescence quenching compared to all test conditions in accordance with its previously characterized GOF properties.

To quantify functional differences, the rate of YFP quenching over the first 60 seconds post NaI addition was calculated from a best-fit line (Fig. 2B). WT BEST1 exhibited a significantly greater quenching rate than both the stuffer control and R218H variant, whereas Y236C displayed a significantly higher quenching rate than WT BEST1 (one-way ANOVA with Dunnett’s multiple-comparisons test, P < 0.0001). Collectively, these results validate the halide-sensitive YFP quenching assay as a robust and quantitative method for assessing the functional impact of BEST1 sequence variants.

### Functional validation of additional *BEST1* variants using the halide-sensitive YFP quenching assay

To further evaluate the ability of the halide-sensitive YFP quenching assay to classify *BEST1* variant function, a panel of nine additional *BEST1* variants with previously reported functional phenotypes was analyzed and compared with published electrophysiological and halide permeability data (Table 1). The selected variants had been characterized in HEK293 cells and/or iPSC-derived retinal pigment epithelium (iPSC-RPE) models using whole-cell patch clamp recordings and/or halide permeability assays.

**Table 1.**
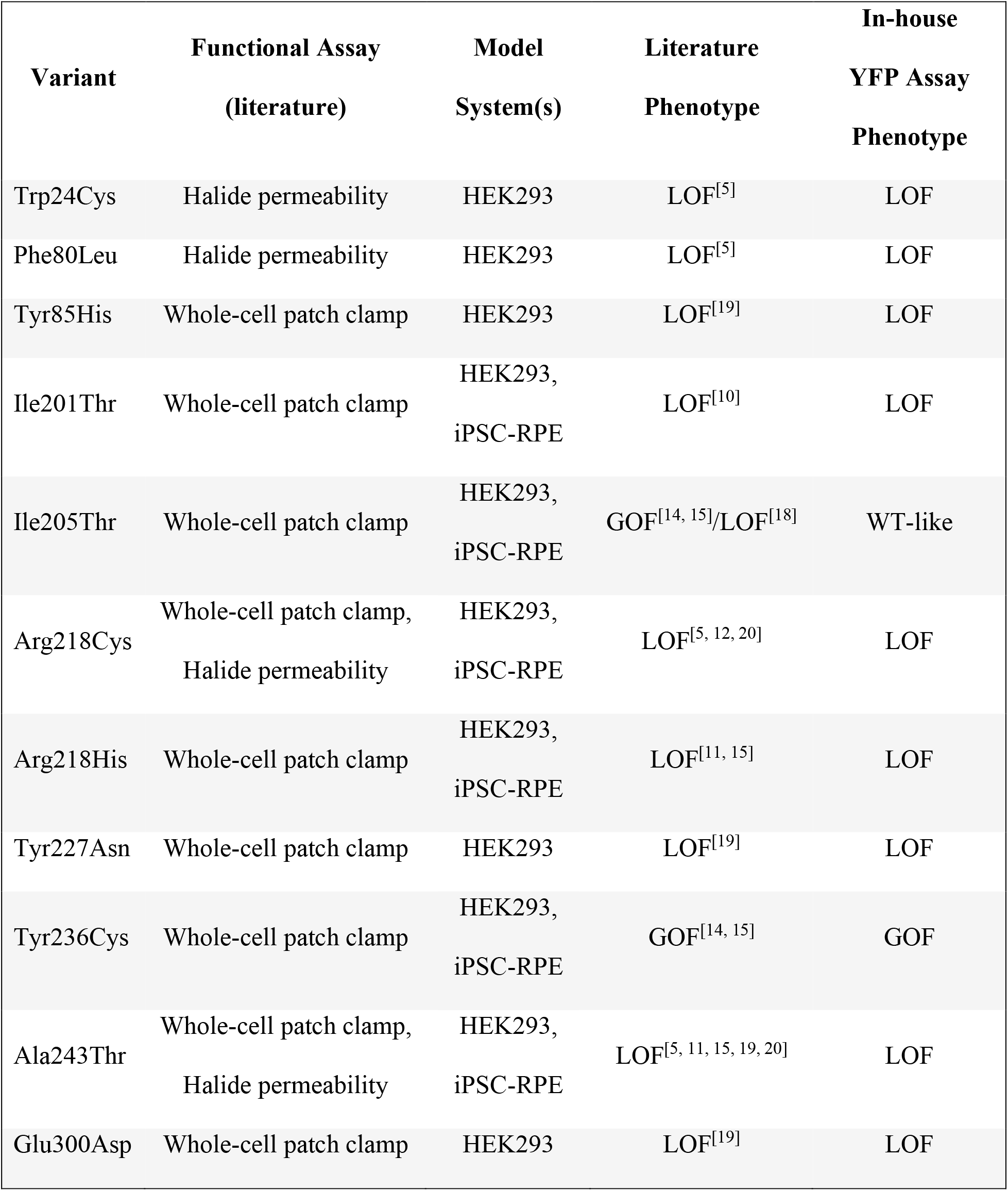
Functional validation of BEST1 variants by halide-sensitive YFP quenching assay relative to published electrophysiological and anion conductance studies. Variants were classified as loss-of-function (LOF), gain-of-function (GOF), or WT-like based on published whole-cell patch clamp and/or halide permeability studies and compared with classifications obtained using the in-house YFP quenching assay. The table includes the well-characterized reference variants R218H (LOF) and Y236C (GOF), which were used for assay validation, together with additional BEST1 variants evaluated in this study. The YFP quenching assay results were highly consistent with literature-reported functional phenotypes across the variants tested.

Trp24Cys, Phe80Leu, Tyr85His, Ile201Thr, Arg218Cys, Tyr227Asn, Ala243Thr, and Glu300Asp exhibited reduced rates of iodide-induced fluorescence quenching relative to WT BEST1 (Supplementary Fig. S3A-H), consistent with loss-of-function phenotypes reported in previous electrophysiological and halide permeability studies.

In contrast, Ile205Thr displayed a quenching rate comparable to WT BEST1 (Supplementary Fig. S3I). Although both gain- and loss-of-function phenotypes have been reported for this variant [14, 15, 18], the results obtained in the YFP quenching assay suggest that Ile205Thr retains channel function under the conditions tested.

Overall, functional classifications obtained using the YFP quenching assay were highly consistent with literature-reported phenotypes derived from whole-cell patch clamp and halide permeability studies. These findings further validate the assay as a reliable and scalable approach for functional characterization of *BEST1* variants and support its utility for distinguishing pathogenic variants with reduced or enhanced channel activity.

## Discussion

Despite the growing number of clinically identified *BEST1* variants, the functional consequences of most mutations remain unknown. As gene therapies for BEST1-associated retinal diseases, including OPGx-BEST1, advance through clinical development, there is an increasing need for high-throughput experimental platforms capable of rapidly distinguishing variants that are likely to respond to gene augmentation therapy from those that may require alternative therapeutic strategies. While electrophysiological studies and patient-derived RPE models have provided important insights into BEST1 biology and disease mechanisms, their low throughput limits systematic evaluation across the expanding catalog of reported variants. In this study, we developed and validated a scalable halide-sensitive YFP quenching assay that provides rapid, quantitative, and reproducible assessment of BEST1 channel activity. Using variants with previously established phenotypes, the assay reliably distinguished loss-of-function and gain-of-function variants and showed strong agreement with published electrophysiological and halide permeability data.

The EYFP-H148Q/I152L reporter used in this study provides a sensitive, indirect measure of BEST1 channel activity through iodide-induced fluorescence quenching. Upon extracellular iodide addition, functional BEST1 channels facilitate iodide influx, resulting in rapid quenching of intracellular YFP fluorescence. Consistent with this mechanism, cells expressing WT BEST1 displayed a significant increase in the rate fluorescence quenching compared with the stuffer control. To validate assay performance, we selected the well-characterized LOF variant R218H and GOF variant Y236C as reference controls. The assay reliably distinguished these variants from WT BEST1, with R218H exhibiting minimal activity comparable to the stuffer control and Y236C demonstrating significantly enhanced channel activity. These findings established the assay’s ability to detect both reduced and increased BEST1 function across a broad dynamic range. Analysis of an additional panel of previously characterized variants further demonstrated strong agreement between YFP quenching assay classifications and reported electrophysiological and halide permeability data, supporting the utility of iodide-induced fluorescence quenching as a surrogate measure of BEST1 channel activity.

The assay measures iodide transport indirectly through fluorescence quenching and therefore does not provide the detailed biophysical information obtainable from electrophysiological recordings. Nevertheless, compared with whole-cell patch clamp electrophysiology, the halide-sensitive YFP assay provides a streamlined workflow, quantitative assessment of channel activity, compatibility with high-throughput screening formats, and the ability to evaluate multiple variants in parallel.

Although the current study evaluated only a limited number of variants, ongoing efforts are expanding the application of this platform to systematically characterize the broader *BEST1* mutational landscape. Comprehensive functional profiling of disease-associated variants will facilitate variant interpretation, inform patient selection for emerging gene therapies such as OPGx-BEST1, and advance our understanding of genotype-phenotype relationships in bestrophinopathies.

In summary, we established a robust and high-throughput platform for functional assessment of BEST1 channel activity. The strong agreement with previously reported functional data supports its application for variant classification and large-scale characterization of disease-associated BEST1 variants, ultimately supporting patient stratification and precision medicine approaches for BEST1-associated retinal disease.

## Materials and methods

### Generation of stable EYFP-H148Q/I152L-expressing HEK293 cells

HEK293 cells (ATCC, Manassas, VA, USA) were transduced with recombinant lentiviral particles encoding the halide-sensitive fluorescent reporter EYFP-H148Q/I152L, which were custom generated by GenScript (Piscataway, NJ, USA). Following puromycin selection, stably transduced cells were expanded and maintained in DMEM supplemented with fetal bovine serum (10% v/v) at 37 °C, 5% CO2. The resulting reporter cell line was used for fluorescence-based assessment of BEST1-mediated halide transport following transient expression of wild-type or mutant BEST1 constructs.

### BEST1 expression constructs

Plasmids encoding wild-type (WT) human BEST1 (NM_004183.4) and disease-associated BEST1 variants were synthesized by VectorBuilder (Chicago, IL, USA) using a custom-designed mammalian expression vector. Plasmid sequences were verified by the manufacturer prior to delivery and were used for transient transfection studies in EYFP-H148Q/I152L-expressing HEK293 reporter cells.

### Cell line transfection

HEK293 cells stably expressing EYFP-H148Q/I152L were seeded in black, clear-bottom, poly-D-lysine-coated 96-well plates (Corning) at a density of 3.5 × 104 cells/well in FluoroBrite™ DMEM supplemented with fetal bovine serum (10% v/v) and 4 mM GlutaMAX™. 24 hours after plating, cells were transiently transfected with 100 ng per well of control vector, wild-type (WT) BEST1, or mutant BEST1 expression plasmids using Lipofectamine™ 3000 (Thermo Fisher Scientific) according to the manufacturer’s instructions. Cells were incubated at 37 °C, 5% CO2 for 48 hours post-transfection prior to functional analysis using the halide-sensitive YFP quenching assay.

### Halide-sensitive YFP quenching assay

48 hours post-transfection, YFP fluorescence was measured using a BioTek Cytation 9 Cell Imaging Microplate Reader (Agilent) using excitation and emission wavelengths of 510 nm and 540 nm, respectively. Baseline fluorescence (F0) was recorded prior to iodide addition. Subsequently, 100 µL of a 2X NaI solution in PBS (without Ca^2+^/Mg^2+^) was dispensed into each well to achieve a final NaI concentration of 75 mM. Changes in YFP fluorescence (F) were then monitored at 7 second intervals for 3 minutes. YFP fluorescence (%) was calculated as F/F0 ×100 for each time point.

### Data analysis

The initial quenching rate was determined from the slope of the normalized fluorescence decay during the first 60 seconds following iodide addition and expressed as % fluorescence/s. Statistical analysis was performed using one-way ANOVA followed by Dunnett’s multiple comparisons test relative to the WT.

