## Supplemental Data for "Functional Validation of Known *BEST1* Variants Using a Robust Halide-Sensitive YFP Quenching Assay"

### **Supplementary figures**

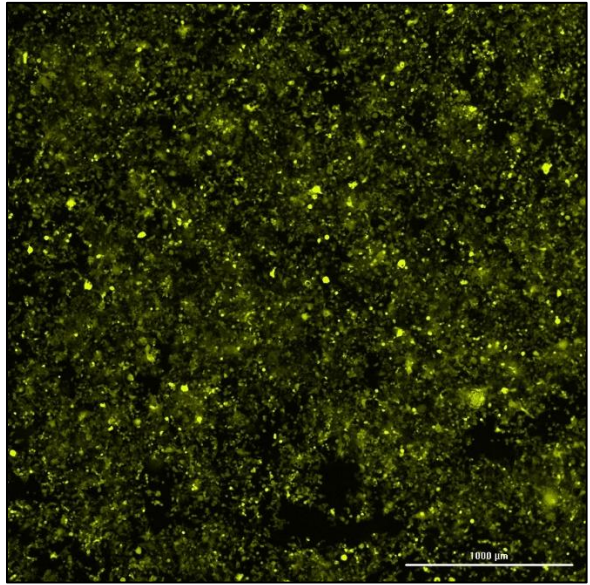

**Supplementary Fig. S1. YFP expression in stable EYFP-H148Q/I152L-expressing HEK293 cells.**

Representative fluorescence micrograph demonstrating robust EYFP expression in the stable reporter cell line used for YFP quenching assay. Scale bar = 1000 $\mu\text{m}$.

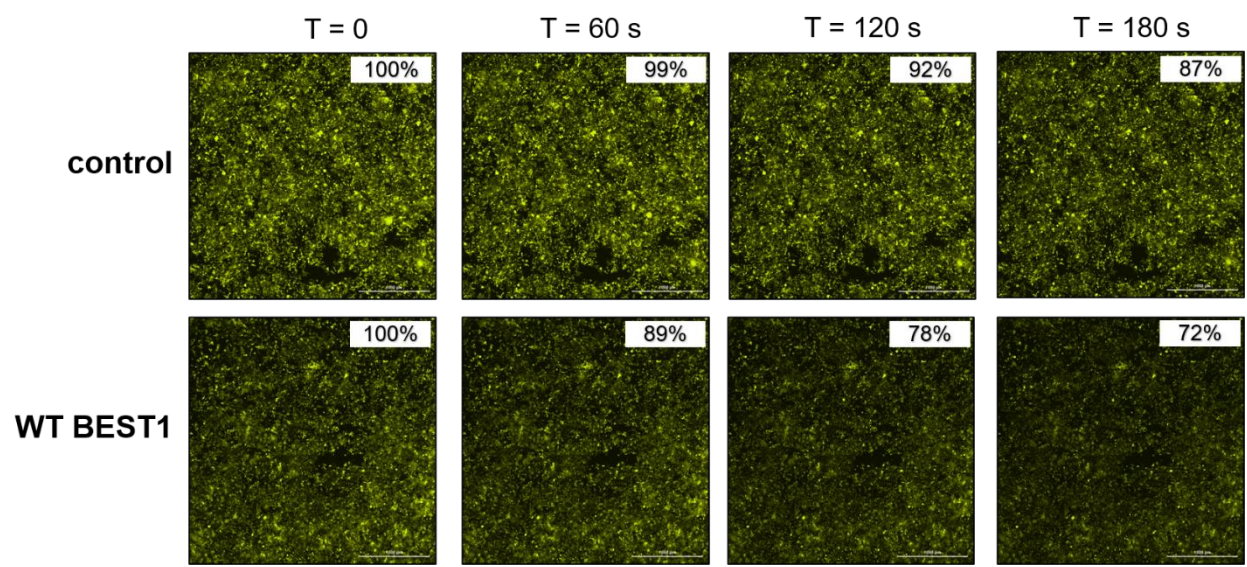

**Supplementary Fig. S2. Representative fluorescence images demonstrating iodide-induced YFP quenching in stable EYFP-H148Q/I152L-expressing HEK293 cells.**

Representative fluorescence images acquired before (0 s) and after extracellular iodide addition (60, 120, and 180 s). Values in the upper right corner of each image indicate fluorescence normalized to the initial fluorescence intensity at 0 s. Cells expressing WT BEST1 exhibit progressive YFP fluorescence quenching following iodide influx, whereas control cells show minimal change. Scale bar = 1000 $\mu\text{m}$.

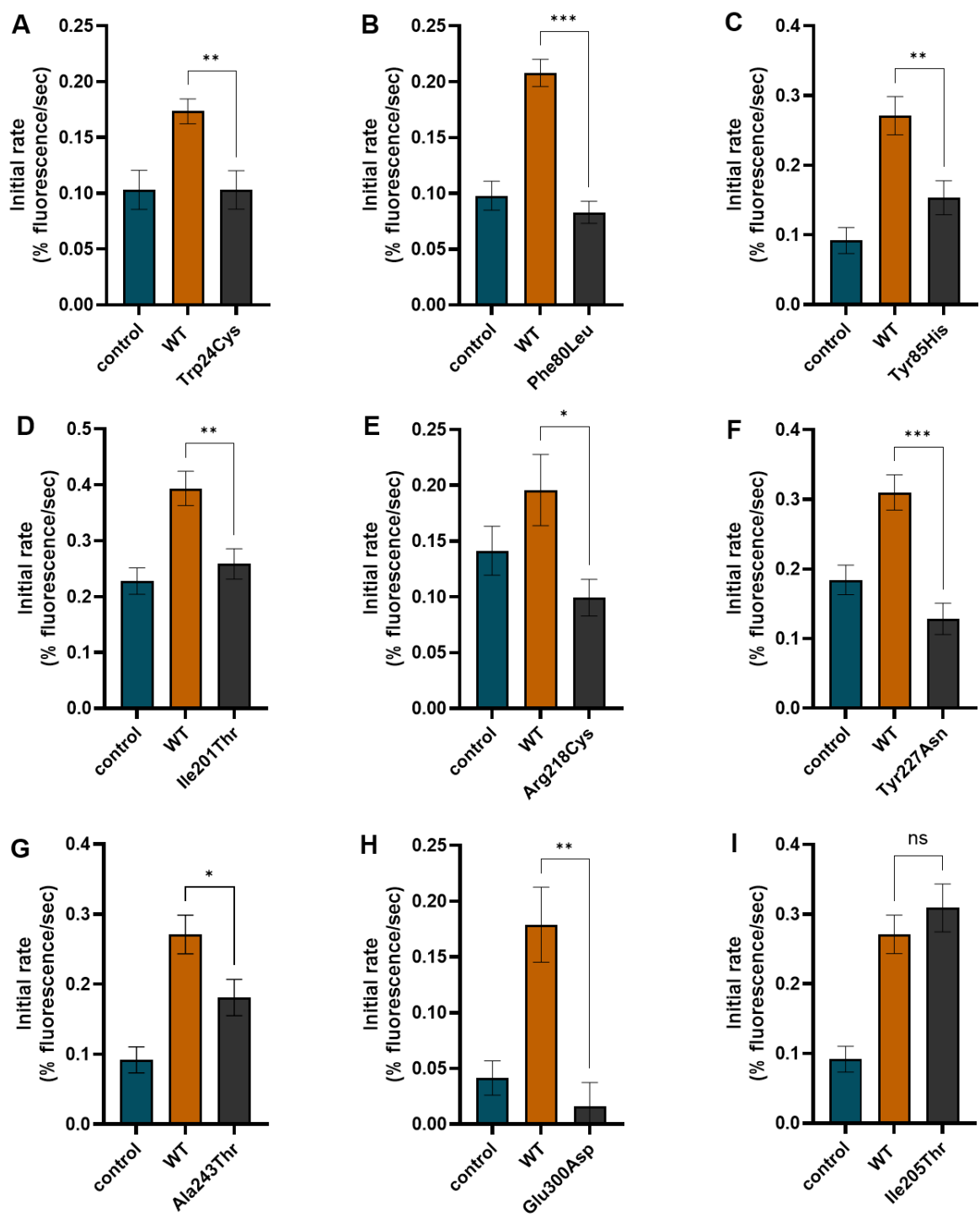

**Supplementary Fig. S3. Initial YFP quenching rates from the in-house YFP quenching assay for additional BEST1 variants previously characterized in electrophysiological and anion conductance studies.**

Initial YFP quenching rates were calculated from fluorescence quenching kinetics during the first 60 s following iodide addition. Statistical significance between WT BEST1 and each mutant was assessed using a two-tailed unpaired Student's t-test with Welch's correction. Data are presented as mean ± SD. \*P < 0.05, \*\*P < 0.01, \*\*\*P < 0.001; ns, not significant.
